# Epigenetic Modifications are Associated with Inter-species Gene Expression Variation in Primates

**DOI:** 10.1101/003467

**Authors:** Xiang Zhou, Carolyn E. Cain, Marsha Myrthil, Noah Lewellen, Katelyn Michelini, Emily R. Davenport, Matthew Stephens, Jonathan K. Pritchard, Yoav Gilad

## Abstract

Changes in gene regulation level have long been thought to play an important role in evolution and speciation, especially in primates. Over the past decade, comparative genomic studies have revealed extensive inter-species differences in gene expression levels yet we know much less about the extent to which regulatory mechanisms differ between species. To begin addressing this gap, we performed a comparative epigenetic study in primate lymphoblastoid cell lines (LCLs), to query the contribution of RNA polymerase II (Pol II) and four histone modifications (H3K4me1, H3K4me3, H3K27ac, and H3K27me3) to inter-species variation in gene expression levels. We found that inter-species differences in mark enrichment near transcription start sites are significantly more often associated with inter-species differences in the corresponding gene expression level than expected by chance alone. Interestingly, we also found that first-order interactions among the histone marks and Pol II do not markedly contribute to the degree of association between the marks and inter-species variation in gene expression levels, suggesting that the marginal effects of the five marks dominate this contribution.

## Introduction

Differences in gene expression level have long been thought to underlie differences in phenotypes between species (Abzhanov et al. 2004; Fay et al. 2004; Shapiro et al. 2004; McGregor et al. 2007), and in particular, to contribute to adaptive evolution in primates (Britten and Davidson 1971; King and Wilson 1975). Consistent with this, previous studies have identified a large number of genes differentially expressed among primates (Enard et al. 2002; Caceres et al. 2003; Karaman et al. 2003; Khaitovich et al. 2004; Khaitovich et al. 2005; Gilad et al. 2006; Blekhman et al. 2008; Blekhman et al. 2009; Babbitt et al. 2010a; Blekhman et al. 2010), and in a few cases, have also found that the interspecies changes in gene expression level might explain differences in complex phenotypes between primates (Rockman et al. 2005; Loisel et al. 2006; Pollard et al. 2006; Prabhakar et al. 2008; Warner et al. 2009; Babbitt et al. 2010b). However, we still know little about the underling regulatory mechanisms leading to the differences in gene expression levels across species. In particular, although a few studies have shown that the inter-species differences in certain epigenetic mechanisms can explain (in a statistical sense) a small proportion of variation in gene expression levels between species (Farcas et al. 2009; Cain et al. 2011; Pai et al. 2011), the relative importance of evolutionary changes in different epigenetic regulatory mechanisms remains largely elusive.

The present study aims to take another step towards understanding gene regulatory evolution in primates, by focusing on inter-species differences in epigenetic regulatory mechanisms that are functionally associated with the regulation of transcription initiation. By studying a number of regulatory mechanisms in parallel in multiple primate species, we can assess the extent to which such differences are associated with inter-species variation in gene expression levels.

We focused on mechanisms associated with transcription initiation, a major determinant of overall steady-state gene expression levels (Wray et al. 2003; Merkin et al. 2012; Tippmann et al. 2012). Transcription of mRNA is preceded by the assembly of large protein complexes that coordinate the recruitment, initiation, and elongation of RNA polymerase II (Pol II) (Woychik and Hampsey 2002). Assembly of these large protein complexes relies on epigenetic information, including various histone modifications (Kouzarides 2007), not only to provide an additional layer of targets for regulatory proteins, but also to directly affect chromatin accessibility of the promoter region to DNA-binding proteins (Felsenfeld and Groudine 2003). As a result, Pol II occupancy and abundance of histone modifications are highly predictive of gene expression levels in multiple cell types (Heintzman et al. 2007; Karlic et al. 2010; Ernst et al. 2011; Bernstein et al. 2012; Tippmann et al. 2012).

A natural hypothesis is that inter-species variation in epigenetic modifications and Pol II abundance could in part contribute to gene expression differences between species. In support of this, a number of examples showed associations between the two. For instance, in *Arabidopsis* leaves, the enrichment of both H3K9ac and H3K4me3 in promoters is associated with transcript abundance between species (Ha et al. 2011). During adipogenesis, orthologous genes with similar expression levels in mouse and human are often marked by similar histone modifications, and orthologous genes associated with inter-species differences in histone modifications are often differentially expressed between species (Mikkelsen et al. 2010). In human, mouse, and pig pluripotent stem cells, the difference in the abundance of several histone modifications correlates with gene expression difference between species (Xiao et al. 2012).

Recent comparative studies of certain epigenetic modifications in primates provide further support for the association between epigenetic modification variation and gene expression variation (Farcas et al. 2009; Cain et al. 2011; Pai et al. 2011; Shulha et al. 2012). For example, Pai et al. showed that inter-species differences in methylation pattern correlate with differences in gene expression level across species (Pai et al. 2011), and Cain et al. found that inter-species differences in the profile of the histone modification H3K4me3 are associated with changes in gene expression level between species (Cain et al. 2011). However, the abundance difference in either of the two marks accounts for only a small proportion of gene expression difference between primates, and it remains unclear whether changes to epigenetic marks play a major causative role in regulatory evolution.

Here, we performed a comparative epigenetic study in primates to query the contribution of Pol II and four histone modifications (H3K4me1, H3K4me3, H3K27ac, and H3K27me3) to inter-species variation in gene expression levels. We choose these five marks not only because their molecular functions have been relatively well studied, but also because they represent a wide variety of transcription initiation regulators. In particular, the four histone modifications mark important regulatory regions: H3K4me1 is present at both active and poised enhancers (Consortium et al. 2007; Heintzman et al. 2007; Koch et al. 2007; Robertson et al. 2008), H3K4me3 marks active transcription start sites (TSSs) (Santos-Rosa et al. 2002; Santos-Rosa et al. 2003; Heintzman et al. 2007; Ruthenburg et al. 2007), H3K27ac marks active enhancers and promoters (Wang et al. 2008; Creyghton et al. 2010; Karlic et al. 2010; Cotney et al. 2012), and H3K27me3 marks repressed genomic regions (Barski et al. 2007; Mikkelsen et al. 2007). In turn, Pol II directly interacts with chromatin remodeling factors (Cho et al. 1998) and catalyzes the transcription of mRNA (Nikolov and Burley 1997).

In what follows, we evaluate the association of each of the five marks with gene expression level variation across species, and further, the joint contribution of all of them to the association with variation in gene expresison, both within, but more importantly between species.

## Results

### Genome-wide profiling of Pol II, four histone marks, and mRNA

We used chromatin immunoprecipitation followed by massively parallel sequencing (ChIPseq) to identify genomic regions associated with Pol II as well as with four histone modifications (H3K4me1, H3K4me3, H3K27ac, and H3K27me3) in lymphoblastoid cell lines (LCLs) from eight individuals from each of the three primate species, humans, chimpanzees, and rhesus macaques (a total of 24 samples for all marks except H3K27ac, for which a rhesus macaque sample is missing; Table S1 and Figure S1). We also extracted RNA from the same 24 LCLs and performed gene expression profiling in each sample by high-throughput sequencing (RNAseq; Table S1 and Figure S1).

As a first step of our analysis we used BWA (Li and Durbin 2009) to align sequence reads to their respective reference genomes (human, hg19; chimpanzee, panTro3; rhesus macaque, rheMac2; Tables S2–S4). Following convention, we then used RSEG (Song and Smith 2011) to identify enriched (broad) regions for H3K27me3 and used MACS (Zhang et al. 2008) to identify (narrow) peaks for the other four marks (Tables S5–S6). To minimize the number of falsely identified mark enrichment differences between species, we used two-step cutoffs to classify the enriched regions/peaks for each mark (Cain et al. 2011). Our approach reflects the assumption that epigenetic profiles in orthologous regions will more often be shared than divergent. Briefly (see Methods for more details), we first used a stringent cutoff to identify enriched regions with high confidence. Conditional on observing an enriched region in one individual using the stringent cutoff, we then classified the same or orthologous regions as enriched in other individuals with a more relaxed second cutoff (Figure S2). Effectively, the more relaxed second threshold borrows information across species to increase power to detect enriched regions in any individual (regardless of species), and reduces the tendency to falsely detect differences in mark abundance between species. Once peak regions were identified, we obtained ‘normalized peak read’ counts for each individual by subtracting the number of mapped reads in the control sample from the number of mapped reads in the ChIPseq sample and further normalizing the resulting values to reads per kilobase per million mapped reads (RPKM) (Mortazavi et al. 2008).

To facilitate comparisons between species of regions that are centered on expressed genes, we used liftOver (Kuhn et al. 2007) to identify orthologous TSSs and followed a previously described approach (Blekhman et al. 2010) to identify orthologous exons. We annotated orthologous TSSs and orthologous exons in a total of 26,115 genes. On the basis of correlations using normalized peak read counts for each of the five marks in regions near the orthologous TSSs and RNA read counts mapped to the orthologous genes, we verified that both the ChIPseq and RNAseq data are of high quality (Figure S3).

### Pol II and four histone modifications are enriched near TSSs

We expected the five marks (Pol II and four histone modifications) to be enriched near TSSs in all three primates, as has been shown previously in other contexts (Barski et al. 2007; Cain et al. 2011; Bernstein et al. 2012; Tippmann et al. 2012; Xiao et al. 2012). To examine this, we considered the average normalized peak read counts in ± 2 kb regions near TSSs across all genes for each individual (more precisely, the regions begin at 2 kb upstream of the TSSs and end at the start of the second orthologous exon or 2 kb downstream of the TSSs, whichever is shorter). Similarly, for each individual, we obtained the normalized peak read counts over the entire genome. We then calculated fold enrichment in regions near TSSs for each mark by considering the ratio of these two values for each individual. We also performed non-parametric Mann-Whitney one-sided tests, based on data from all eight individuals in each species, to determine whether the normalized peak read counts in TSS regions are significantly higher than their genome-wide counterparts. The results of these analyses clearly indicate that all five marks are significantly enriched near TSSs, regardless of species (Figure 1A). The enrichment pattern is robust with respect to the choice of the size of the TSS region, but gradually decreases for increasingly larger regions around TSSs (Figure S4).

**Figure 1.**
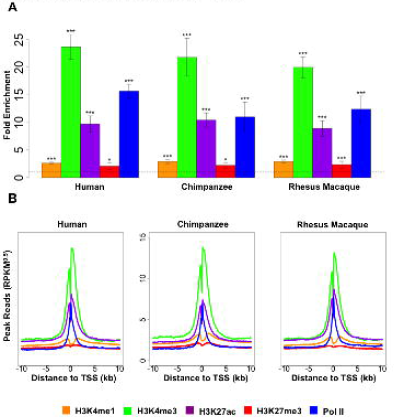
Marks are enriched near TSSs. (A) Fold enrichment of the five marks in ± 2 kb regions near TSSs in the three primates. Error bars indicate standard deviation calculated across eight individuals in each species. Asterisks indicate significance levels based on Mann-Whitney one-sided tests (* *p*<0.05, ** *p*<0.01, *** *p*<0.001). (B) Distribution of normalized peak read counts for five marks around TSSs for each of the three primates. Units are in square root of RPKM (i.e. RPKM^0.5^) and are averaged across individuals and across genes.

To explore the localization pattern of the five marks near TSSs, we generated, for each species, the distributions of normalized peak read counts averaged across all genes and all individuals (Figure 1B). Consistent with previous studies (Barski et al. 2007; Heintzman et al. 2007; Schones et al. 2008; Cain et al. 2011; Bernstein et al. 2012; Tippmann et al. 2012; Xiao et al. 2012), all five marks display bimodal distribution patterns near TSSs – albeit to a lesser extent for H3K27me3 – with two modes flanking the TSSs.

Levels of the five marks are also highly correlated with each other in regions near TSSs (Figure S5). Specifically, H3K27me3 levels are negatively correlated with the other four marks, while H3K4me1, H3K4me3, H3K27ac and Pol II levels are positively correlated with each other.

### Mark abundance near TSSs correlates with gene expression levels within species

To explore the relationship between mark abundance and gene expression levels, we first obtained quantitative measurements and performed appropriate transformations for both mark enrichment level and RNA expression level (see Methods for details). Next, we divided genes evenly (thus, arbitrarily) into the following three sets based on their expression levels: highly expressed, intermediately expressed and expressed at low levels. We obtained the distribution of the mark enrichment levels near TSSs, averaged across individuals within a species and across genes in each given set (Figures 2A, S6A, and S7A). Regardless of species, we found that the repressive mark H3K27me3 (Barski et al. 2007; Mikkelsen et al. 2007) is enriched near TSSs of genes expressed at low levels, whereas Pol II and the other four active histone marks (Nikolov and Burley 1997; Santos-Rosa et al. 2002; Santos-Rosa et al. 2003; Consortium et al. 2007; Heintzman et al. 2007; Koch et al. 2007; Ruthenburg et al. 2007; Robertson et al. 2008; Wang et al. 2008; Creyghton et al. 2010; Karlic et al. 2010; Cotney et al. 2012) are highly enriched near TSSs of highly expressed genes. To verify that these patterns are robust, we arbitrarily divided genes into a larger number of groups based on absolute gene expression levels, such that each group contains 200 genes (except the first group, which contains all non-expressed genes, and the last group, which contains fewer than 200 genes). We plotted the mean mark enrichment levels in the ± 2 kb region near TSSs against the mean gene expression levels in each group, both averaged across individuals within a species and across genes in that group (Figures 2B, S6B, and S7B). We again observed a negative trend between the enrichment levels of H3K27me3 and gene expression levels, as well as positive trends for the correlations between the enrichment levels of the other four marks and gene expression levels. These trends were robust with respect to the choice of TSS region size (Figure S8).

**Figure 2.**
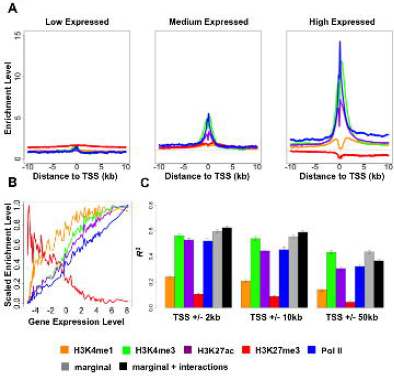
Mark enrichment levels are correlated with gene expression levels in human. (A) Density of enrichment level for five marks around TSSs for genes with low, medium, and high expression levels. Values are averaged across individuals and across genes in each category. (B) Mark enrichment levels plotted against gene expression levels for sliding windows of genes (n=200) ordered from low to high expression levels. Enrichment levels are obtained in ± 2 kb regions near TSSs and scaled to be between 0 and 1. All values are averaged across individuals and across genes in the window. (C) Proportion of variance in gene expression levels explained (R squared) by individual marginal effects (five colored bars), combined mark marginal effects (grey bars) and all first-order interaction effects in addition to marginal effects (black bars) of the five marks. Results are shown for enrichment levels in TSS regions with increasing length. Error bars indicate standard deviation calculated based on 20 split replicates.

To quantitatively measure the relationship, namely the extent of association, between mark abundance and gene expression levels across genes within each species, we fitted a linear model with gene expression level as response and mark enrichment level in regions near TSSs as covariates (averaged across individuals). In addition, to avoid model over-fitting, we used a 10-fold cross-validation (with 20 split replicates) and calculated R squared, in the test set (Figures 2C, S6C, S7C, and S9). We found that the R squared by H3K4me3, H3K27ac, or Pol II is much higher than the R squared by the other two marks. Our observations with respect to individual marks are in close agreement with results from previous studies in other tissues (Karlic et al. 2010; Dong et al. 2012; Tippmann et al. 2012). In a statistical sense, levels of the five marks combined explain approximately 58% of the variance in gene expression levels within species (59% in human, 58% in chimpanzee, and 57% in rhesus macaque).

Because the marks show strong correlation patterns near TSSs (Figure S5) and because previous studies have shown that combinatorial patterns of histone modifications and Pol II (i.e. chromatin states) could be of biological importance (Ernst and Kellis 2010; Ernst et al. 2011), we asked if adding interaction effects increases the R squared. To do so, we considered all first-order interactions among marks – including all interactions between two marks, among three marks, etc. – in addition to their marginal effects. We used a Bayesian variable selection regression (BVSR) model (Mitchell and Beauchamp 1988; George and Mcculloch 1993; Guan and Stephens 2011; Zhou and Stephens 2012) with gene expression level as response and all marginal and interaction terms as covariates. BVSR provides a “posterior inclusion probability” (PIP) for each covariate, which indicates the confidence that the covariate contributes to prediction of phenotype. In addition, BVSR can produce reliable estimates of the proportion of variance explained by all covariates (Guan and Stephens 2011; Zhou and Stephens 2012). We used the posterior means as coefficient estimates and calculated R squared in the test set (Figures 2C, S6C, and S7C). Using this approach, we found that all marginal effects, except for H3K4me1, are important features that are consistently selected by the model (PIP > 0.9; Figure S10). Among the interaction features, interactions H3K27ac-Pol II, H3K4me3-H3K27ac with or without Pol II, H3K4me1-H3K27ac with or without Pol II, H3K4me1-H3K4me3 with or without Pol II, H3K4me1-H3K27me3 with or without H3K4me3, are consistently selected as important features (PIP > 0.9; Figure S10).

Somewhat surprisingly, considering all interaction features does not increase much the association of the marks with variation in gene expression levels across genes within species (black bars versus grey bars, Figures 2C, S6C, and S7C).

### Differences in mark enrichment are associated with gene expression differences across species

Next, we considered differences between species. As a first step, we identified differentially expressed (DE) genes across species, as well as orthologous TSS regions that are associated with interspecies differences in enrichment of histone marks or Pol II. As expected, we found a smaller number of differences between humans and chimpanzees, than between either humans or chimpanzees, and rhesus macaques (Tables 1, S7, and S8).

**Table 1.** Number of TSS regions associated with interspecies differences in enrichment of marks and number of DE genes from pairwise comparisons among three primates at an FDR cutoff of 5%.

|  | **H3K4me1** | **H3K4me3** | **H3K27ac** | **H3K27me3** | **Pol II** | **RNA** |
| --- | --- | --- | --- | --- | --- | --- |
| H vs C | 137 | 3037 | 3176 | 438 | 1577 | 3824 |
| H vs R | 3298 | 5257 | 5549 | 1487 | 3708 | 6567 |
| C vs R | 3421 | 4928 | 5456 | 1017 | 3299 | 5914 |

We found that DE genes, compared with non-DE genes, are more likely to show inter-species differences in mark enrichment at the TSSs (Figures 3A). The directions of the associations are consistent with our expectations (namely, we observed increased gene expression associated with decrease in H3K27me3 and increase in the other marks and Pol II). In addition, for those genes where the mark enrichment levels and the gene expression levels differ in the expected direction between species (i.e. opposite direction for H3K27me3, same direction for the other four marks), DE genes are generally more often associated with inter-species differences in mark enrichment at their TSS regions than expected by chance alone (Figure 3B). These observations are robust with respect to the choice of false discovery rate (FDR) cutoff for classifying DE genes (Figure S11).

**Figure 3.**
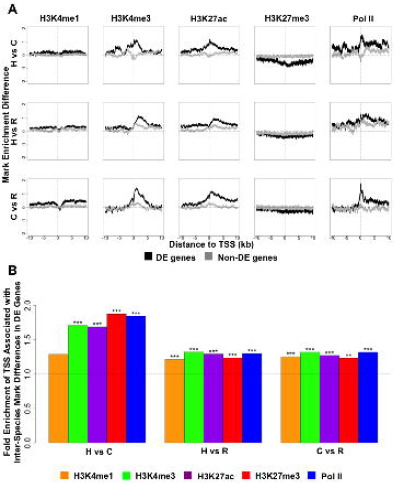
DE genes associate with inter-species differences in mark enrichment at TSSs. (A) Enrichment level differences for the five marks around TSSs of DE genes (black) and non-DE genes (grey) for each pair of species. Mark differences are considered with respect to the species associated with the lower gene expression level. DE genes are determined based on an FDR cutoff of 5%. (B) TSS regions associated with inter-species differences in any mark are enriched for DE genes. Plotted is the fold enrichment of TSS regions associated with inter-species differences in enriched marks in DE genes across pairs of species, for genes where the mark enrichment levels and the gene expression levels differ in the expected direction (i.e. opposite for H3K27me3, same for the other four marks). Both the TSS regions associated with inter-species differences in enriched marks and DE genes are determined based on an FDR cutoff of 5%. Asterisks indicate significance levels from binomial tests (* *p*<0.05, ** *p*<0.01, *** *p*<0.001). H: human; C: chimpanzee; R: rhesus macaque.

The association of inter-species DE genes and differences in mark enrichment in the corresponding TSS regions across species encouraged us to further explore this relationship. We performed regression analyses similar to those described above, except that we focused on differences in gene expression level and mark enrichment level between pairs of species.

Considering data from each pair of species at a time (e.g., human and chimpanzee), we divided genes into 200-gene groups based on inter-species expression level difference and plotted the mean mark enrichment level differences against the mean gene expression level differences across the species (Figure 4A). We found that differences in mark enrichment level correlate with differences in gene expression level between primates. In particular, the difference in H3K27me3 enrichment level is negatively correlated with gene expression level differences between species, and the enrichment level differences of the other four marks are positively correlated with inter-species DE. A few representative patterns are shown in Figure S12. These observations are robust with respect to the chosen size of the TSS regions (Figure S13).

**Figure 4.**
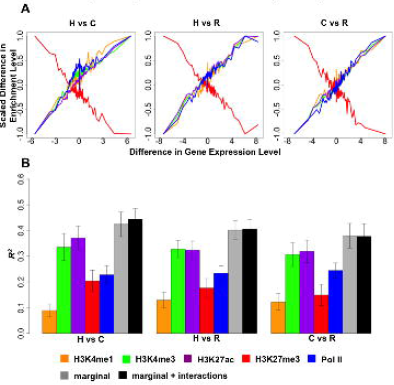
Differences in mark enrichment level correlate with differences in gene expression level between pairs of primates. (A) Differences in mark enrichment level is plotted against differences in gene expression level for sliding windows of genes (n=200) ordered from based on the DE effect size. Differences in enrichment level are obtained in ± 2 kb regions near TSSs and scaled to be between −1 and 1. All values are averaged across individuals and across genes in the window. (B) Proportion of variance in gene expression level differences explained (R squared) by mark enrichment level differences, for all pairwise comparisons among the three primates. Different linear models are fitted to account for individual marginal effects (five colored bars), combined marginal effects (grey bars) and all first-order interaction effects in addition to marginal effects (black bars) of the five marks. The DE genes are determined based on an FDR cutoff of 5%. Enrichment level differences are obtained in ± 2 kb regions. Error bars indicate standard deviation calculated across 20 split replicates. H: human; C: chimpanzee; R: rhesus macaque.

To quantitatively measure the proportion of variance in inter-species gene expression level differences explained by the five marks, either individually or combined, we again used a 10-fold cross-validation strategy and applied linear models to calculate R squared in DE genes (Figures 4B, S14, and S15). We focused on the ± 2 kb regions near TSSs as we found these to be most predictive in the analysis of data within species. Each of the five marks explained an appreciable proportion of variance in gene expression level differences between any pairs of species (Figure 4B). The relative importance of the five marks is consistent with that observed within species (Figures 4B and 2C). Together, the five marks explain (in a statistical sense) approximately 40% of the variance in gene expression levels across species (42% between human and chimpanzee, 40% between human and rhesus macaque, and 38% between chimpanzee and rhesus macaque; FDR < 5%).

Finally, we used BVSR to select important marginal and first-order interaction features (Figures 4B, 5 and S14). Again, we found that all marginal effects are important features that are consistently selected by the model (PIP > 0.9 for all FDR cutoffs; Figure 5). However, no interaction term is consistently selected as important feature for pairs of species across a range of FDR cutoffs. In addition, modeling the interaction features in addition to the marginal effects does not increase the overall explained variance in gene expression level differences between primates (Figures 4B and S14).

**Figure 5.**
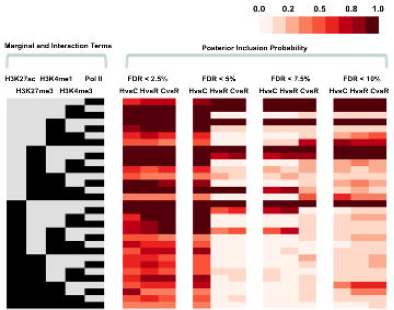
Importance of marginal and first-order interaction effects from five marks for explaining gene expression level differences between primates. The left panel lists all interaction terms among the five marks, where each row represents an interaction term and each column represents the presence (black) or absence (grey) of a particular mark effect for that interaction term. For example, the first row represents the marginal effect of Pol II, and the seventh row represents the interaction effect of H3K4me1, H3K4me3, and Pol II. The right panel lists the corresponding posterior inclusion probability of each term in the BVSR between any pairs of primates, for DE genes classified with different FDR cutoffs. The posterior inclusion probability measures the importance of each interaction term, with values ranging between 0 and 1; higher values indicate more importance. Mark enrichment level differences ± 2 kb regions near TSSs are used for fitting. H: human; C: chimpanzee; R: rhesus macaque.

## Discussion

We explored the extent to which inter-species differences in Pol II and four histone modifications are associated with differences in gene expression levels across primates. We found that all five marks combined explain 40% of the variation in gene expression levels between pairs of species (when we focused on DE genes), which is 5% more than the single most informative mark. These observations suggest that epigenetic modifications are substantially associated with changes in gene expression level among primates and may represent important molecular mechanisms in primate evolution.

### Correlation and causality

As we briefly mention in the results section, it is important to clarify that we use the words “contribute” and “explain” to mean a purely statistical conditional relationship between the mark abundance and gene expression levels.

Previous work that focused on molecular mechanisms indicates that variation in Pol II and histone modifications directly affect gene regulation. Specifically, it is well established that Pol II directly transcribes mRNA (Nikolov and Burley 1997). It has been shown that H3K4me3 recruits chromatin-remodeling complexes to increase the accessibility of the chromatin to transcriptional machinery and therefore promote gene expression (Mizuguchi et al. 1997; Santos-Rosa et al. 2003; Ruthenburg et al. 2007). It is also generally believed that the other three histone modifications (H3K4me1, H3K27ac, H3K27me3) act in a similar fashion as H3K4me3 to either promote or inhibit gene expression by regulating chromatin accessibility (Felsenfeld and Groudine 2003). In particular, the clearance of H3K4me1 is shown to be necessary for the subsequent binding of some transcription factors (Lupien et al. 2008).

On the other hand, recent work (from our lab as well) indicates that oftentimes differences in histone marks are mediated by changes in transcription factor binding (Kasowski et al. 2013; Kilpinen et al. 2013; McVicker et al. 2013). Transcription factor binding may be the principle determinant of chromatin state, which is then stabilized or marked by histone modifications. In that sense, the association between changes in histone modification across species and variation in gene expression levels may indicate a direct causal relationship, but rather an indirect one, possibly mediated by inter-species differences in transcription factor binding.

Indeed, we did not perform experiments here that allow us to directly infer causality. The well-established links from previous studies imply that the quantitative relationship between mark abundance and gene expression level likely reflect, at least in part, a (direct or indirect) causal contribution. Indeed, the larger R squared by H3K4me3, H3K27ac and Pol II compared with the other two marks is consistent with the key functions of the three in promoting transcription (Mizuguchi et al. 1997; Cho et al. 1998; Santos-Rosa et al. 2003; Ruthenburg et al. 2007; Chen et al. 2011). Nevertheless, we caution against the over-interpretation of these association results, and defer the interrogation of both the direct and directional effects of epigenetic marks on gene expression levels to future studies. It is also possible that other molecular mechanisms are responsible for the correlation between mark abundance and gene expression levels, at least for a subset of the marks and in a subset of the genes. For example, in some cases a true causal factor may independently affect both gene expression level and histone modifications at the same location (this has been demonstrated previously in other contexts (Rybtsova et al. 2007; Edmunds et al. 2008)), causing correlations between the two. Our study was not designed to distinguish between all of these possible scenarios.

Regardless of whether the abundance of the four histone modifications and Pol II are truly causally related to variation in gene expression levels, they are only involved in some of the many intermediate steps that a complex machinery takes to convert genome sequence variation, including both *cis-* and trans-acting sequence differences, into gene expression variation. The amount of gene expression variation explained by the five marks, therefore, at best, still reflects only part of the causal contribution of the sequence variation to gene expression variation through transcriptional processes (as opposed to other aspects of the mRNA life cycle, such as decay). In any case, the proportion of variance in gene expression levels tracing back to the sequence variation through the five marks is likely smaller than what we have observed here (because the mark abundance variation is at a later step than the sequence variation). If the abundance levels of the five marks are not causal but are by-products of some true causal factors (such as variation in transcription factor binding), then the proportion of variance in gene expression levels tracing back to the sequence variation through these true causal factors could be larger than what we have observed here (because the mark abundance levels are noisy measurements of these causal factors). Moreover, the effects from the sequence variation could be in complicated forms, because simple measurements of sequence conservation and sequence divergence do not predict gene expression level difference between species (Figure S16). It will be of great interest to reveal the detailed steps of this process and the ultimate contribution of sequence variation to gene expression variation by mapping all the different regulatory checkpoints.

### The chain of events

In our work, we followed the example of previous studies (Ernst and Kellis 2010; Tippmann et al. 2012) and treated the abundance of Pol II and histone modifications equivalently in investigating their relationship to gene expression level variation. We note that numerous studies have established a direct role of Pol II in transcription initiation while pointing to indirect roles of the four histone modifications in transcription initiation through Pol II (Mizuguchi et al. 1997; Nikolov and Burley 1997; Cho et al. 1998; Felsenfeld and Groudine 2003; Santos-Rosa et al. 2003; Ruthenburg et al. 2007). These observations suggest that it might make sense to apply a two-stage analysis to the data. First, we might investigate the contribution of the four histone modifications to Pol II abundance (Figures S17A and S17C), and then investigate the contribution of Pol II abundance to gene expression levels (Figures 2C and 4B). However, such naïve analyses ignore the contribution of the four histone modifications to gene expression levels through mechanisms other than regulating the recruitment of Pol II and its *abundance* levels. For example, studies have shown that Pol II abundance itself is not the sole determinant of transcription initiation, and Pol II can remain in a pausing state without initiating active transcription (Guenther et al. 2007; Zeitlinger et al. 2007; Core and Lis 2008; Core et al. 2008). Such a pausing state can be predicted by histone modifications (Chen et al. 2011). Indeed, in the present study, we showed that modeling the five marks together explains a higher proportion of variation in gene expression level than would be explained by Pol II alone (Figures 2C and Figure 4B). In fact, for both within-species and inter-species analysis, the R squared by the four histone modifications is only slightly smaller than that by the four histone modifications and Pol II (Figures S17B and S17D). In addition, the PIPs for each interaction term among the four histone modifications are not sensitive to whether Pol II is included in the analysis or not (i.e. the PIPs for each interaction term analyzed without Pol II are similar to those obtained by first analyzing with Pol II but then marginalizing out Pol II; data not shown). As a result of these considerations, we chose to treat the abundance of Pol II and histone modifications equivalently in our study.

### The contribution of interactions between marks

In addition to the marginal effects of the five marks, we also explored the importance of all first-order interaction effects among them. In particular, we identified several notable interaction effects that are important to explaining (in a statistical sense) gene expression level variation within species. Many of these effects are present in important chromatin states identified by other computational methods (Ernst and Kellis 2010; Ernst et al. 2011). Two of these interactions, one between H3K4me1 and H3K27ac, and the other between H3K4me1 and H3K27me3, have been recognized to be part of important classes of genomic elements during early development in humans (Rada-Iglesias et al. 2011). However, we found it surprising that the explained proportion of variance in gene expression levels (within or between species) remains largely similar, whether or not we consider all first-order interactions in addition to the marginal effects in the model. Our results imply that the marginal effects of the five marks dominate the contribution and interaction effects contribute only a small proportion.

It is possible that we are underpowered to identify important interactions. Indeed, measurement noise for any interaction effect is likely the multiplication of noise levels accompanying each marginal effect, and in the case of the inter-species analysis, the sample size is small (because we focused on differentially expressed genes). In addition, in both the within and between species analyses, we assessed only interaction effects for marks near TSSs. It therefore remains possible that interactions among marks in distal but specific regions (e.g. enhancer regions) may affect gene expression levels to a larger extent. The statistical challenges and consideration of a narrow region near the TSSs notwithstanding, the lack of important and consistent interaction effects in our data is nevertheless an intriguing observation.

### Using LCLs as a model system

In the present study, we chose to work with LCLs because they provide abundant material and represent a homogenous cell type from all three species. We note that using LCLs have been criticized previously for two main reasons: that LCLs are cultured cells instead of a primary tissue and are susceptible to batch effects (Akey et al. 2007; Choy et al. 2008), and that LCLs require an initial virus transformation that may causes artifacts (Hannula et al. 2001; Carter et al. 2002; Redon et al. 2006). However, numerous previous studies have demonstrated the usefulness of LCLs in genomics studies (Monks et al. 2004; Morley et al. 2004; Cheung et al. 2005; Stranger et al. 2005; Dixon et al. 2007; International HapMap et al. 2007; Moffatt et al. 2007; Stranger et al. 2007; Veyrieras et al. 2008; Ge et al. 2009), and have shown that the regulatory architectures identified in LCLs are highly replicable in primary tissues (Bullaughey et al. 2009; Dimas et al. 2009; Verlaan et al. 2009; Ding et al. 2010; Zeller et al. 2010). In particular, it has been shown that the patterns of inter-species gene expression level differences in LCLs highly resemble those in primary tissues between primates (Khaitovich et al. 2006). In the present study, we also found that the contribution of the five marks to gene expression level variation within species highly resembles those obtained in other tissues or organisms (Karlic et al. 2010; Dong et al. 2012; Tippmann et al. 2012), suggesting that a similar quantitative relationship between the five marks and gene expression level variation exists across multiple species and tissues. In addition, the number of DE genes detected from LCLs in the present study is similar to that obtained from liver tissue in a different study (Blekhman et al. 2010), and an average of 28% of the DE genes from our study are also identified as DE genes in theirs (20% between human and chimpanzee, 33% between human and rhesus macaque, and 31% between chimpanzee and rhesus macaque; FDR < 5%). Therefore, although we acknowledge the potential pitfalls of using LCLs, we believe that they provide a useful and reasonable system, and that the genomic mechanisms we interrogated in LCLs are likely representative of those in primary tissues.

### Final remarks

Even if we assume direct or indirect causality, we note that Pol II and all four histone modifications together do not explain all intra- or inter-species gene expression level variation. Indeed, even with an overly simplified model that accounts for noise in mark enrichment measurement or gene expression measurement (see Methods for details), the “maximal contribution” from the five marks together to gene expression variation is still estimated to be only 59% within species (60% for human, 59% for chimpanzee and 58% for rhesus macaque), and 43% for DE genes between species (47% between human and chimpanzee, 43% between human and rhesus macaque, and 40% between chimpanzee and rhesus macaque; FDR < 5%). It is likely that other molecular mechanisms (for example, those affecting transcription initiation and RNA decay) account for the remaining portion of variation in gene expression levels. We hope that, by collecting comparative genomic data on additional epigenetic and genetic regulatory mechanisms, obtaining more accurate measurements and furthering our analysis on various interactions in the future, we could eventually obtain a better understanding of the detailed molecular mechanisms underlying the evolution of gene expression levels in primates.

## Materials and Methods

### Samples and cell culture

Eight LCLs from human, chimpanzee, and rhesus macaque individuals were obtained from Coriell Institute (http://www.coriell.org/), New Iberia Research Center (University of Louisiana at Lafayette), and New England Primate Research Center (NEPRC, Harvard Medical School). In addition, one input sample from each of the three species was used as control. Cell lines were grown at 37°C in RPMI media with 15% FBS, supplemented with 2 mM L-glutamate, 100 I.U./mL penicillin, and 100 μg/mL streptomycin.

### ChIPseq and RNAseq

Chromatin immunoprecipitation (ChIP) was performed largely as previously described (Cain et al. 2011). In addition to the data collected in this study, we incorporated data from 6 H3K4me3 ChIP assays performed in one previous study (Cain et al. 2011) and 5 Pol II ChIP assays performed in another (Pai et al. 2012). For newer samples that were not described in these two previous studies, chromatin was sheared with a Covaris S2 (settings: 40 minutes, duty cycle 20%, intensity 8, 200 cycles/burst, 500uL at a time in 12 x 24mm tubes). The amount of antibody used for each ChIP was separately optimized for H3K4me3 (4 ug, Abcam ab8580), H3K4me1 (12 ug, Millipore 07-436), H3K27ac (4 ug, Abcam ab4729), H3K27me3 (4 ug, Millipore 07-449), and Pol II (10 ug, Santa Cruz sc-9001). Part of the data for the human samples is also used in another study (McVicker et al. 2013).

The quality of each immunoprecipitation was assessed by RT-PCR of positive and negative control genomic regions previously shown to be enriched or not enriched in ENCODE LCL ChIP data for each feature (Consortium et al. 2011). Successful ChIP assays showed enrichment at the positive control regions relative to the negative control regions in the immunoprecipitated sample compared to the input whole-cell extract from the same individual. We prepared Illumina sequencing libraries from the DNA from each ChIP sample, and from a pooled input sample from each species (containing equal amounts of DNA by mass from each individual in a species) as previously described (Marioni et al. 2008), starting with 20 uL of ChIP output or 4 ng pooled input sample.

Libraries were sequenced in one or more lanes on an Illumina sequencing system using standard Illumina protocols. H3K4me1, H3K4me3, H3K27ac, and H3K27me3 samples were sequenced on a Genome Analyzer II (GAII) system (single end, 36 bp), and Pol II and input samples were sequenced on a HiSeq system (single end, 28 bp and 50 bp, respectively). Input reads were trimmed to 28 bp and 36 bp, where appropriate, for comparison to the reads generated from ChIP samples.

For RNAseq, RNA was extracted and processed to create Illumina sequencing libraries as previously described (Marioni et al. 2008; Cain et al. 2011). Each sample was sequenced on one or more lanes of an Illumina GAII system.

### Reads alignment

All sequenced reads were aligned to human (hg19, February 2009), chimpanzee (panTro3, October 2010), or rhesus macaque (rheMac2, January 2006) genome builds with BWA (Li and Durbin 2009) version 0.5.9. Each genome was slightly modified to exclude the Y chromosome, mitochondrial DNA, and regions labeled as random.

We excluded ChIPseq and input reads that were assigned a quality score less than 10, contain more than 2 mismatches or any gaps compared to the reference genome, or are duplicates. We excluded RNAseq reads that were assigned a quality score less than 10 or contain more than 2 mismatches or any gaps relative to the reference genome.

### Classifying genomic regions as enriched

MACS version 1.4.1 (Zhang et al. 2008) was used to identify sharp peaks of enrichment for H3K4me1, H3K4me3, H3K27ac, and Pol II; RSEG version 0.4.4 (Song and Smith 2011) was used to classify enrichment of broad genomic regions of enrichment for H3K27me3. For MACS, we specified an initial *p* value threshold that was optimized for each feature (H3K4me1, 0.01; H3K4me3, 0.0001; H3K27ac, 0.001; and Pol II, 0.001), with the appropriate species’ input control file for comparison. Because the chimpanzee sequenced input sample yielded roughly twice the number of reads as the other input samples, to avoid any species-bias related to number of input reads, we subsampled the chimpanzee input data to a final number of 40 million reads, which is now comparable to the human and rhesus macaque input samples. For RSEG, we used the “rseg-diff’ function with input control data, with the recommended 20 maximum iterations for hidden Markov model training.

Enriched regions or peaks identified by MACS or RSEG were next filtered to exclude regions or peaks that could not be mapped uniquely in all three primate genomes. To do so, we first divided the genome into 200 bp windows, and we retained those windows that could be mapped to all three primate genomes with gaps less than 100 bp using liftOver (Kuhn et al. 2007), and that have at least 80% of bases mappable across all three species (where mappability was measured by the ability of 20 bp sequences to be uniquely mapped to a genome). We then excluded enriched regions or peaks that did not overlap this set of 200 bp windows. To further ensure that regions or peaks of enrichment for features have orthologous positions in human, chimpanzee, and rhesus macaque genomes, we also mapped each region or peak coordinates to the other two genomes with liftOver and excluded enriched regions and peaks that failed to map with at least 20% of the bases aligning to the other genomes.

To minimize the number of falsely identified differences in enrichment status between individuals, we applied two-step cutoffs (Cain et al. 2011) to classify enriched regions or peaks for each mark. (We chose to present data with this two-step cutoffs procedure because this procedure was also used in other stages of the analysis, though the results presented here are not very sensitive to whether this procedure is applied.) Specifically, for the features analyzed with MACS, we chose a first, stringent FDR cutoff based on the distributions of FDR values associated with identified peaks. A first cutoff of 5% FDR was chosen because we observe a clear enrichment below that value for all features. To select the more relaxed cutoff, we examined the distributions of FDR values for peaks overlapping orthologous positions of peaks that pass the first cutoff (where the orthologous regions were classified by liftOver. These distributions are enriched for small values, which is consistent with individuals of the same or a closely related species having similar epigenetic profiles. We chose secondary FDR cutoffs to capture this enrichment for each feature (H3K4me1, 15%; H3K4me3, 10%; H3K27ac, 15%; and Pol II, 10%).

For H3K27me3, which was analyzed with RSEG, we could not choose cutoffs exactly the same way as described above because RSEG does not produces an FDR value for each enriched region. Instead, for each region classified as enriched, RSEG assigns a domain score, which is the sum of the posterior scores of all bins within the domain. To choose a first, stringent score cutoff, we calculated the proportion of regions classified as enriched by RSEG that overlap regions classified as enriched in ENCODE LCL data (Consortium et al. 2011) at a range of score cutoffs. We chose a first, stringent, score cutoff of 20 because ∼85% of regions classified as enriched with a score of at least 20 overlapped regions classified as enriched in ENCODE data. To choose a second, more relaxed, score cutoff, we examined all the regions classified as enriched that overlap the orthologous positions of regions classified as enriched by the first cutoff. As expected, over 80% of these regions overlap ENCODE enriched regions, consistent with a low rate of false-positive calls of enrichment among this set of regions. We therefore chose the second, more relaxed cutoff for enrichment to be classification as enriched by RSEG, without a score requirement.

### Mark enrichment level and RNA expression level

We mapped RNA sequencing reads to each orthologous exon, summed values across exons for each gene, and normalized them with respect to the total mapped reads and total exon length to obtain the normalized reads (in RPKM) for each gene. Following convention (Ouyang et al. 2009; Dong et al. 2012; Tippmann et al. 2012), we transformed these normalized reads by log2 transformation (after adding a small value to ensure positive values (Ouyang et al. 2009; Dong et al. 2012)), and we termed the resulting value *gene expression level*. For the five marks, we divided the number of normalized peak reads in different sized regions surrounding the TSSs for each gene by the genome-wide average to obtain mark fold enrichment in these regions. We performed square root transformation following previous studies (Pique-Regi et al. 2011), and termed the resulting value *mark enrichment level*, which serves as a measurement of mark abundance. We note that the normalized peak read counts require a step to subtract reads in the corresponding region from input controls, but the final results presented here are not sensitive to whether this step is performed or not.

### Analysis with Bayesian variable selection regression models

BVSR specifies sparse priors on covariates, and has been proven to be effective in selecting important features as well as to be accurate in estimating the proportion of variance in phenotypes explained by all covariates (Guan and Stephens 2011; Zhou and Stephens 2012). To fit BVSR, we first standardized each covariate to have unit standard deviation. We then used the Markov chain Monte Carlo method (10000 burn-in iterations and 100000 sampling iterations) to obtain posterior samples of parameters, using the software piMASS with default settings (Guan and Stephens 2011). For R squared estimation, we fitted the model in the training set and used the posterior means as coefficient estimates to calculate R squared in the test set. For PIP calculation, we fitted the model using both training and test sets.

### Classifying DE genes and TSS regions associated with inter-species differences in mark enrichment

We tested all genes whose median mark enrichment level or gene expression level across 16 individuals in the species being compared is above zero. To ensure that values are comparable across individuals, we first quantile transformed either the gene expression level or the mark enrichment level across genes in each individual into a standard normal distribution. Afterwards, to guard against model misspecification, for each gene, we further quantile transformed either the gene expression level or the mark enrichment level (in the +/− 2 kb region near the TSSs) in 16 individuals from the two species being compared into a standard normal distribution. We then fitted a linear model in these individuals with sex as a covariate and species label as a predictor. We tested whether the coefficient for the species label is significantly different from zero. At the same time, we constructed a null distribution by permuting every possible combination of the species label (a total of 6435 combinations for H3K27ac and 12870 combinations for the other four marks and RNA) and we calculated the FDR based on this empirical null.

### Overlap between DE genes and TSS regions associated with inter-species differences in mark enrichment

For each mark, we focused on genes where the gene expression levels and mark enrichment levels differ between pairs of species in the expected direction. Specifically, for H3K27me3, we focused on genes where the inter-species gene expression level and the mark enrichment level differences are in the opposite direction. For other four marks, we focused on genes where the interspecies gene expression level and the mark enrichment level differences are in the same direction. Afterwards, we divided the proportion of DE genes that also have TSS regions that are associated with inter-species differences in mark enrichment, by the proportion of non-DE genes that have TSS regions that are associated with inter-species differences in mark enrichment, in order to calculate fold enrichment. We used the binomial test to obtain the corresponding *p* values.

### Measuring sequence conservation and difference between species

We used four different measurements for sequence conservation as well as sequence difference between pairs of species in the TSS region. To measure sequence conservation, we obtained the average Phastcons score (Siepel et al. 2005) and the PhyloP score (Cooper et al. 2005; Pollard et al. 2010) in the TSS region. To measure sequence difference, we first used blastn to obtain a list of aligned sequences between pairs of species. We then calculated the proportion of aligned sequence in the TSS region between pairs of species as one measurement, and calculated the average percentage of identity in these aligned sequence in the TSS as another measurement.

### Estimating “maximal” R squared by accounting for measurement noise

Here, we estimate the “maximal” R squared by the five marks, by taking into account the measurement noise accompanying both mark enrichment levels and gene expression levels. We consider the following linear model:

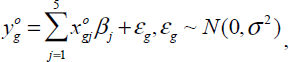

where 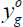 is the observed phenotype (i.e. gene expression level or gene expression level difference, averaged across individuals) for the *g*th gene, 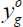 is the observed *j*th covariate (i.e. enrichment level or enrichment level difference for *j*th mark, averaged across individuals) for the *g*th gene, *ε*_*g*_ is the error term, which follows a normal distribution with variance *σ*^2^. For convenience, we assume that both phenotypes and covariates are already mean centered.

We assume that both 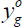 and 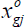 are noisy measurements of the true underlying phenotype *y*_*g*_ and covariate *x*_*gj*_, with the corresponding noises following independent normal distributions

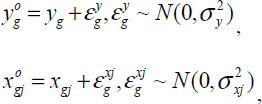

where 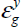 and 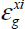 are assumed to be independent across genes and independent of each other.

With the above assumptions, we have

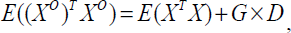

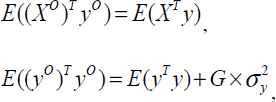

where *G* is the number of genes, *X*^*o*^ is a *G* by *5* matrix with *gj*th element 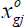, *X* is a *G* by *5* matrix with gjth element *x*_*gj*_, *y*^*o*^ is a G-vector with gth element 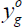, *y* is a *G*-vector with *g*th element *y*_*g*_, and 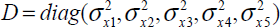 is a diagonal matrix.

Therefore, we can approximate the “maximal” R squared by

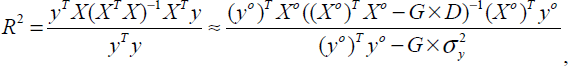

and we replace 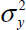 and 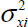 with the estimated values

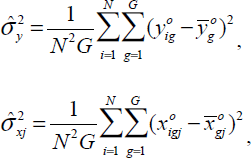

where *N* is the number of individuals.

## Acknowledgments

We thank the New England Primate Research Center, the New Iberia Research Center, and the Yerkes primate center for primate LCLs. We thank Ran Blekhman for providing a list of orthologous exons, Roger Pique-Regi for assistance in identifying orthologous TSSs, Jacob Degner and Graham McVicker for read mapping assistance, and Timothee Flutre, Ester Pantaleo, Dessilava Petkova and Heejung Shim for helpful comments on the manuscript. We thank all members of the Gilad, Pritchard and Stephens labs for insightful discussions. This was supported by NIH grants GM077959 and GM084996 to YG. The University of Louisiana at Lafayette New Iberia Research Center is funded by National Institutes of Health/National Center for Research Resources (NIH/NCRR) grants RR015087, RR014491, and RR016483, and the Genetics Core of the New England Primate Research Center by NIH/NCRR grant RR00168.

---

Figure S1. An illustration of the study design.

Figure S2. Choices of cutoffs for classifying regions as enriched. (A-H) Histograms of peaks of H3K4me3 (A-B), H3K4me1 (C-D), H3K27ac (E-F), and Pol II (G-H) enrichment, as classified by MACS, at various FDR thresholds. (A, C, E, and G) All peaks with FDR ≤ 50%; the dashed line indicates the stringent 5% cutoff. (B, D, F, and H) Peaks with FDR ≤ 50% that overlap a peak with FDR ≤ 5% in another individual; the relaxed FDR cutoff for each feature is marked by a dashed line. (I-J) Number of H3K27me3 enriched regions (dark squares, left axis) as classified by RSEG, and the proportion of those regions overlapping ENCODE H3K27me3 peaks (light triangles, right axis) at various score cutoffs up to 200. (I) All enriched regions; the dashed line indicates the stringent 20 score cutoff. (J) Enriched regions that overlap enriched regions with ≥ 20 score from another individual; the relaxed score cutoff is 0 – that is, any region classified as “enriched” by RSEG.

Figure S3. Pairwise Spearman’s rank correlations between individuals from the three primates for four histone marks, Pol II and RNA. Calculations are based on mark abundance in ± 2 kb regions near orthologous TSSs for five marks, and on gene expression level in orthologous exons for RNA. H: human; C: chimpanzee; R: rhesus macaque.

Figure S4. Fold enrichment of the five marks in ± 10 kb (A) and ± 50 kb (B) regions near TSSs in three primates. Error bars indicate standard deviation calculated across all genes and all individuals. Asterisks indicate significance levels (* *p*<0.05, ** *p*<0.01, *** *p*<0.001).

Figure S5. Pairwise Spearman’s rank correlations between marks in each of the three primates. Calculations are based on mark abundance in ± 2 kb (A), ± 10 kb (B), ± 50 kb (C) regions near orthologous TSSs.

Figure S6. Mark enrichment levels are correlated with gene expression levels in chimpanzee. Legends are identical to those in Figure 2.

Figure S7. Mark enrichment levels are correlated with gene expression levels in rhesus macaque. Legends are identical to those in Figure 2.

Figure S8. Mark enrichment levels are plotted against gene expression levels for sliding windows of genes (n=200) ordered by increasing expression levels in the three primates. Enrichment levels are obtained in either ± 10 kb or ± 50 kb regions near TSSs and scaled to be between 0 and 1. All values are averaged across individuals and across genes in the window.

Figure S9. Scatterplot of predicted gene expression levels against true gene expression levels for all analyzed genes in human. Predicted values are obtained based on linear models with either individual marginal effects (colored plots) or all marginal mark effects (grey plot) using mark enrichment levels in ± 2 kb regions near TSSs.

Figure S10. Importance of the marginal and first-order interaction effects from the five marks for explaining gene expression levels in the three primates. The left panel lists all interaction terms among the five marks; each row represents an interaction term and each column represents the presence (black) or absence (grey) of a particular mark effect for that interaction term. For example, the first row represents the marginal effect of Pol II, and the seventh row represents the interaction effect of H3K4me1, H3K4me3 and Pol II. The right panel lists the corresponding posterior inclusion probability of each term in the BVSR in the three species. The posterior inclusion probability measures the importance of each interaction term, with values ranging between 0 and 1; higher values indicate more importance. Mark enrichment levels ± 2 kb regions near TSSs are used for fitting. H: human; C: chimpanzee; R: rhesus macaque.

Figure S11. TSS regions associated with inter-species differences in enriched marks are enriched for differentially expressed (DE) genes. TSS regions associated with inter-species differences in enriched marks and DE genes are determined by various FDR cutoffs (2.5%, 7.5%, and 10%). Legends are identical to those in Figure 3. H: human; C: chimpanzee; R: rhesus macaque.

Figure S12. An example of mark abundance and gene expression levels across three species. The x-axis is the distance along a genomic region containing the gene REX02. The y-axes show RNAseq reads (black), as well as ChIPseq reads for the five marks (color) and input controls (grey), all scaled with respect to the total mapped read counts.

Figure S13. Differences in mark enrichment level plotted against differences in gene expression level for sliding windows (n=200) of genes ordered by increasing differences in expression level. Differences in enrichment level are obtained in either ± 10kb or ± 50kb regions near TSSs and scaled to be between −1 and 1. All values are averaged across individuals and across genes in the window. H: human; C: chimpanzee; R: rhesus macaque.

Figure S14. Proportion of variance in gene expression level differences explained (R squared) by mark enrichment level differences, for all pairwise comparisons among the three primates. Different linear models are fitted to account for individual effects (five colored bars), combined marginal effects (grey bars) and all first-order interaction effects in addition to marginal effects (black bars) of the five marks. DE genes are determined based on an FDR cutoff of 5%. Enrichment level differences are obtained in ± 2 kb regions. Error bars indicate standard deviation calculated across 20 split replicates. H: human; C: chimpanzee; R: rhesus macaque.

Figure S15. Scatterplot of predicted gene expression level differences plotted against true gene expression level differences for DE genes between human and chimpanzee. Predicted values are obtained based on linear models using either individual mark effects (colored plots) or all marginal mark effects (grey plot) with mark enrichment level differences in ± 2 kb regions near TSSs. DE genes are determined based on an FDR cutoff of 5%. H: human; C: chimpanzee; R: rhesus macaque.

Figure S16. Scatterplot of gene expression level differences plotted against sequence conservation and sequence divergence between pairs of species. Two sequence conservation measurement and two sequence divergence measurements are used. DE genes are determined based on an FDR cutoff of 5%. H: human; C: chimpanzee; R: rhesus macaque.

Figure S17. R squared by four histone modifications and Pol II or by four histone modifications alone. (A) Proportion of variance in Pol II enrichment level explained by enrichment level of histone modifications. (B) Proportion of variance in gene expression level explained by mark enrichment level. (C) Proportion of variance in Pol II enrichment level differences explained by enrichment level differences of histone modifications in DE genes. (D) Proportion of variance in gene expression level differences explained by mark enrichment level differences in DE genes. Different linear models are fitted to account for combined marginal effects (grey bars) and all first-order interaction effects in addition to marginal effects (black bars). DE genes are determined based on an FDR cutoff of 5%. Enrichment level differences are obtained in ± 2 kb regions. Error bars indicate standard deviation calculated across 20 split replicates. H: human; C: chimpanzee; R: rhesus macaque.

Table S1. Characteristics and sources of lymphoblastoid cell lines.

Table S2. Number of total sequenced reads for each feature for each individual.

Table S3. Number of total mapped reads with quality score >10 for each feature for each individual.

Table S4. Number of sequenced and mapped reads for pooled input samples.

Table S5. Number of enriched regions/peaks identified for each feature for each individual.

Table S6. Number of mapped reads in enriched regions/peaks for each mark for each individual.

Table S7. Number of tested TSS regions and genes.

Table S8. Number of TSS regions associated with inter-species differences in enriched marks and number of differentially expressed genes identified at different FDR cutoff.

## Reference

Abzhanov A, Protas M, Grant BR, Grant PR, Tabin CJ. 2004. Bmp4 and morphological variation of beaks in Darwin's finches. Science 305(5689): 1462–1465.

Akey JM, Biswas S, Leek JT, Storey JD. 2007. On the design and analysis of gene expression studies in human populations. Nature genetics 39(7): 807–808; author reply 808–809.

Babbitt CC, Fedrigo O, Pfefferle AD, Boyle AP, Horvath JE, Furey TS, Wray GA. 2010a. Both noncoding and protein-coding RNAs contribute to gene expression evolution in the primate brain. Genome Biol Evol 2: 67–79.

Babbitt CC, Silverman JS, Haygood R, Reininga JM, Rockman MV, Wray GA. 2010b. Multiple Functional Variants in cis Modulate PDYN Expression. Mol Biol Evol 27(2): 465–479.

Barski A, Cuddapah S, Cui K, Roh TY, Schones DE, Wang Z, Wei G, Chepelev I, Zhao K. 2007. High-resolution profiling of histone methylations in the human genome. Cell 129(4): 823–837.

Bernstein BE, Birney E, Dunham I, Green ED, Gunter C, Snyder M. 2012. An integrated encyclopedia of DNA elements in the human genome. Nature 489(7414): 57–74.

Blekhman R, Marioni JC, Zumbo P, Stephens M, Gilad Y. 2010. Sex-specific and lineage-specific alternative splicing in primates. Genome Res 20(2): 180–189.

Blekhman R, Oshlack A, Chabot AE, Smyth GK, Gilad Y. 2008. Gene regulation in primates evolves under tissue-specific selection pressures. PLoS Genet 4(11): e1000271.

Blekhman R, Oshlack A, Gilad Y. 2009. Segmental duplications contribute to gene expression differences between humans and chimpanzees. Genetics 182(2): 627–630.

Britten RJ, Davidson EH. 1971. Repetitive and non-repetitive DNA sequences and a speculation on the origins of evolutionary novelty. Q Rev Biol 46(2): 111–138.

Bullaughey K, Chavarria CI, Coop G, Gilad Y. 2009. Expression quantitative trait loci detected in cell lines are often present in primary tissues. Hum Mol Genet 18(22): 4296–4303.

Caceres M, Lachuer J, Zapala MA, Redmond JC, Kudo L, Geschwind DH, Lockhart DJ, Preuss TM, Barlow C. 2003. Elevated gene expression levels distinguish human from non-human primate brains. Proc Natl Acad Sci USA 100(22): 13030–13035.

Cain CE, Blekhman R, Marioni JC, Gilad Y. 2011. Gene expression differences among primates are associated with changes in a histone epigenetic modification. Genetics 187(4): 1225–1234.

Carter KL, Cahir-McFarland E, Kieff E. 2002. Epstein-Barr virus-induced changes in B-lymphocyte gene expression. J Virol 76(20): 10427–10436.

Chen Y, Jorgensen M, Kolde R, Zhao X, Parker B, Valen E, Wen J, Sandelin A. 2011. Prediction of RNA Polymerase II recruitment, elongation and stalling from histone modification data. BMC genomics 12: 544.

Cheung VG, Spielman RS, Ewens KG, Weber TM, Morley M, Burdick JT. 2005. Mapping determinants of human gene expression by regional and genome-wide association. Nature 437(7063): 1365–1369.

Cho H, Orphanides G, Sun X, Yang XJ, Ogryzko V, Lees E, Nakatani Y, Reinberg D. 1998. A human RNA polymerase II complex containing factors that modify chromatin structure. Mol Cell Biol 18(9): 5355–5363.

Choy E, Yelensky R, Bonakdar S, Plenge RM, Saxena R, De Jager PL, Shaw SY, Wolfish CS, Slavik JM, Cotsapas C et al. 2008. Genetic analysis of human traits in vitro: drug response and gene expression in lymphoblastoid cell lines. PLoS genetics 4(11): e1000287.

Consortium EP, Birney E, Stamatoyannopoulos JA, Dutta A, Guigo R, Gingeras TR, Margulies EH, Weng Z Snyder M Dermitzakis ET et al. 2007. Identification and analysis of functional elements in 1% of the human genome by the ENCODE pilot project. Nature 447(7146): 799–816.

Consortium EP, Myers RM, Stamatoyannopoulos J, Snyder M, Dunham I, Hardison RC, Bernstein BE, Gingeras TR, Kent WJ, Birney E et al. 2011. A user’s guide to the encyclopedia of DNA elements (ENCODE). PLoS biology 9(4): e1001046.

Cooper GM, Stone EA, Asimenos G, Green ED, Batzoglou S, Sidow A. 2005. Distribution and intensity of constraint in mammalian genomic sequence. Genome Res 15(7): 901–913.

Core LJ, Lis JT. 2008. Transcription regulation through promoter-proximal pausing of RNA polymerase II. Science 319(5871): 1791–1792.

Core LJ, Waterfall JJ, Lis JT. 2008. Nascent RNA sequencing reveals widespread pausing and divergent initiation at human promoters. Science 322(5909): 1845–1848.

Cotney J, Leng J, Oh S, Demare LE, Reilly SK, Gerstein MB, Noonan JP. 2012. Chromatin state signatures associated with tissue-specific gene expression and enhancer activity in the embryonic limb. Genome research.

Creyghton MP, Cheng AW, Welstead GG, Kooistra T, Carey BW, Steine EJ, Hanna J, Lodato MA, Frampton GM, Sharp PA et al. 2010. Histone H3K27ac separates active from poised enhancers and predicts developmental state. Proceedings of the National Academy of Sciences of the United States of America 107(50): 21931–21936.

Dimas AS, Deutsch S, Stranger BE, Montgomery SB, Borel C, Attar-Cohen H, Ingle C, Beazley C, Gutierrez Arcelus M, Sekowska M et al. 2009. Common regulatory variation impacts gene expression in a cell type-dependent manner. Science 325(5945): 1246–1250.

Ding J, Gudjonsson JE, Liang L, Stuart PE, Li Y, Chen W, Weichenthal M, Ellinghaus E, Franke A, Cookson W et al. 2010. Gene expression in skin and lymphoblastoid cells: Refined statistical method reveals extensive overlap in cis-eQTL signals. Am J Hum Genet 87(6): 779–789.

Dixon AL, Liang L, Moffatt MF, Chen W, Heath S, Wong KC, Taylor J, Burnett E, Gut I, Farrall M et al. 2007. A genome-wide association study of global gene expression. Nature genetics 39(10): 1202–1207.

Dong X, Greven MC, Kundaje A, Djebali S, Brown JB, Cheng C, Gingeras TR, Gerstein M, Guigo R, Birney E et al. 2012. Modeling gene expression using chromatin features in various cellular contexts. Genome biology 13(9): R53.

Edmunds JW, Mahadevan LC, Clayton AL. 2008. Dynamic histone H3 methylation during gene induction: HYPB/Setd2 mediates all H3K36 trimethylation. EMBOJ 27(2): 406–420.

Enard W, Khaitovich P, Klose J, Zollner S, Heissig F, Giavalisco P, Nieselt-Struwe K, Muchmore E, Varki A, Ravid R et al. 2002. Intra- and interspecific variation in primate gene expression patterns. Science 296(5566): 340–343.

Ernst J, Kellis M. 2010. Discovery and characterization of chromatin states for systematic annotation of the human genome. Nat Biotechnol 28(8): 817–825.

Ernst J, Kheradpour P, Mikkelsen TS, Shoresh N, Ward LD, Epstein CB, Zhang X, Wang L, Issner R, Coyne M et al. 2011. Mapping and analysis of chromatin state dynamics in nine human cell types. Nature 473(7345): 43–49.

Farcas R, Schneider E, Frauenknecht K, Kondova I, Bontrop R, Bohl J, Navarro B, Metzler M, Zischler H, Zechner U et al. 2009. Differences in DNA methylation patterns and expression of the CCRK gene in human and nonhuman primate cortices. Mol Biol Evol 26(6): 1379–1389.

Fay JC, McCullough HL, Sniegowski PD, Eisen MB. 2004. Population genetic variation in gene expression is associated with phenotypic variation in Saccharomyces cerevisiae. Genome Biol 5(4): R26.

Felsenfeld G, Groudine M. 2003. Controlling the double helix. Nature 421(6921): 448–453.

Ge B, Pokholok DK, Kwan T, Grundberg E, Morcos L, Verlaan DJ, Le J, Koka V, Lam KC, Gagne V et al. 2009. Global patterns of cis variation in human cells revealed by high-density allelic expression analysis. Nature genetics 41(11): 1216–1222.

George EI, Mcculloch RE. 1993. Variable Selection Via Gibbs Sampling. J Am Stat Assoc 88(423): 881–889.

Gilad Y, Oshlack A, Smyth GK, Speed TP, White KP. 2006. Expression profiling in primates reveals a rapid evolution of human transcription factors. Nature 440(7081): 242–245.

Guan YT, Stephens M. 2011. Bayesian Variable Selection Regression for Genome-Wide Association Studies and Other Large-Scale Problems. Ann Appl Stat 5(3): 1780–1815.

Guenther MG, Levine SS, Boyer LA, Jaenisch R, Young RA. 2007. A chromatin landmark and transcription initiation at most promoters in human cells. Cell 130(1): 77–88.

Ha M, Ng DW, Li WH, Chen ZJ. 2011. Coordinated histone modifications are associated with gene expression variation within and between species. Genome research 21(4): 590–598.

Hannula K, Lipsanen-Nyman M, Scherer SW, Holmberg C, Hoglund P, Kere J. 2001. Maternal and paternal chromosomes 7 show differential methylation of many genes in lymphoblast DNA. Genomics 73(1): 1–9.

Heintzman ND, Stuart RK, Hon G, Fu Y, Ching CW, Hawkins RD, Barrera LO, Van Calcar S, Qu C, Ching KA et al. 2007. Distinct and predictive chromatin signatures of transcriptional promoters and enhancers in the human genome. Nature genetics 39(3): 311–318.

International HapMap C Frazer KA Ballinger DG Cox DR Hinds DA Stuve LL Gibbs RA Belmont JW Boudreau A Hardenbol P et al. 2007. A second generation human haplotype map of over 3.1 million SNPs. Nature 449(7164): 851–861.

Karaman MW, Houck ML, Chemnick LG, Nagpal S, Chawannakul D, Sudano D, Pike BL, Ho VV, Ryder OA, Hacia JG. 2003. Comparative analysis of gene-expression patterns in human and African great ape cultured fibroblasts. Genome Res 13(7): 1619–1630.

Karlic R, Chung HR, Lasserre J, Vlahovicek K, Vingron M. 2010. Histone modification levels are predictive for gene expression. Proceedings of the National Academy of Sciences of the United States of America 107(7): 2926–2931.

Kasowski M, Kyriazopoulou-Panagiotopoulou S, Grubert F, Zaugg JB, Kundaje A, Liu Y, Boyle AP, Zhang QC, Zakharia F, Spacek DV et al. 2013. Extensive variation in chromatin states across humans. Science 342(6159): 750–752.

Khaitovich P, Enard W, Lachmann M, Paabo S. 2006. Evolution of primate gene expression. Nat Rev Genet 7(9): 693–702.

Khaitovich P, Hellmann I, Enard W, Nowick K, Leinweber M, Franz H, Weiss G, Lachmann M, Paabo S. 2005. Parallel patterns of evolution in the genomes and transcriptomes of humans and chimpanzees. Science 309(5742): 1850–1854.

Khaitovich P, Muetzel B, She X, Lachmann M, Hellmann I, Dietzsch J, Steigele S, Do HH, Weiss G, Enard W et al. 2004. Regional patterns of gene expression in human and chimpanzee brains. Genome Res 14(8): 1462–1473.

Kilpinen H, Waszak SM, Gschwind AR, Raghav SK, Witwicki RM, Orioli A, Migliavacca E, Wiederkehr M, Gutierrez-Arcelus M, Panousis NI et al. 2013. Coordinated effects of sequence variation on DNA binding, chromatin structure, and transcription. Science 342(6159): 744–747.

King MC, Wilson AC. 1975. Evolution at two levels in humans and chimpanzees. Science 188(4184): 107–116.

Koch CM, Andrews RM, Flicek P, Dillon SC, Karaoz U, Clelland GK, Wilcox S, Beare DM, Fowler JC, Couttet P et al. 2007. The landscape of histone modifications across 1% of the human genome in five human cell lines. Genome research 17(6): 691–707.

Kouzarides T. 2007. Chromatin modifications and their function. Cell 128(4): 693–705.

Kuhn RM, Karolchik D, Zweig AS, Trumbower H, Thomas DJ, Thakkapallayil A, Sugnet CW, Stanke M, Smith KE, Siepel A et al. 2007. The UCSC genome browser database: update 2007. Nucleic acids research 35(Database issue): D668–673.

Li H, Durbin R. 2009. Fast and accurate short read alignment with Burrows-Wheeler transform. Bioinformatics 25(14): 1754–1760.

Loisel DA, Rockman MV, Wray GA, Altmann J, Alberts SC. 2006. Ancient polymorphism and functional variation in the primate MHC-DQA1 5′ cis-regulatory region. Proceedings of the National Academy of Sciences of the United States of America 103(44): 16331–16336.

Lupien M, Eeckhoute J, Meyer CA, Wang Q, Zhang Y, Li W, Carroll JS, Liu XS, Brown M. 2008. FoxA1 translates epigenetic signatures into enhancer-driven lineage-specific transcription. Cell 132(6): 958–970.

Marioni JC, Mason CE, Mane SM, Stephens M, Gilad Y. 2008. RNA-seq: an assessment of technical reproducibility and comparison with gene expression arrays. Genome Res 18(9): 1509–1517.

McGregor AP, Orgogozo V, Delon I, Zanet J, Srinivasan DG, Payre F, Stern DL. 2007. Morphological evolution through multiple cis-regulatory mutations at a single gene. Nature 448(7153): 587–590.

McVicker G, van de Geijn B, Degner JF, Cain CE, Banovich NE, Raj A, Lewellen N, Myrthil M, Gilad Y, Pritchard JK. 2013. Identification of genetic variants that affect histone modifications in human cells. Science 342(6159): 747–749.

Merkin J, Russell C, Chen P, Burge CB. 2012. Evolutionary dynamics of gene and isoform regulation in Mammalian tissues. Science 338(6114): 1593–1599.

Mikkelsen TS, Ku M, Jaffe DB, Issac B, Lieberman E, Giannoukos G, Alvarez P, Brockman W, Kim TK, Koche RP et al. 2007. Genome-wide maps of chromatin state in pluripotent and lineage-committed cells. Nature 448(7153): 553–560.

Mikkelsen TS, Xu Z, Zhang X, Wang L, Gimble JM, Lander ES, Rosen ED. 2010. Comparative epigenomic analysis of murine and human adipogenesis. Cell 143(1): 156–169.

Mitchell TJ, Beauchamp JJ. 1988. Bayesian Variable Selection in Linear-Regression. J Am Stat Assoc 83(404): 1023–1032.

Mizuguchi G, Tsukiyama T, Wisniewski J, Wu C. 1997. Role of nucleosome remodeling factor NURF in transcriptional activation of chromatin. Mol Cell 1(1): 141–150.

Moffatt MF, Kabesch M, Liang L, Dixon AL, Strachan D, Heath S, Depner M, von Berg A, Bufe A, Rietschel E et al. 2007. Genetic variants regulating ORMDL3 expression contribute to the risk of childhood asthma. Nature 448(7152): 470–473.

Monks SA, Leonardson A, Zhu H, Cundiff P, Pietrusiak P, Edwards S, Phillips JW, Sachs A, Schadt EE. 2004. Genetic inheritance of gene expression in human cell lines. Am J Hum Genet 75(6): 1094–1105.

Morley M, Molony CM, Weber TM, Devlin JL, Ewens KG, Spielman RS, Cheung VG. 2004. Genetic analysis of genome-wide variation in human gene expression. Nature 430(7001): 743–747.

Mortazavi A, Williams BA, McCue K, Schaeffer L, Wold B. 2008. Mapping and quantifying mammalian transcriptomes by RNA-Seq. Nat Methods 5(7): 621–628.

Nikolov DB, Burley SK. 1997. RNA polymerase II transcription initiation: a structural view. Proceedings of the National Academy of Sciences of the United States of America 94(1): 15–22.

Ouyang Z, Zhou Q, Wong WH. 2009. ChIP-Seq of transcription factors predicts absolute and differential gene expression in embryonic stem cells. Proceedings of the National Academy of Sciences of the United States of America 106(51): 21521–21526.

Pai AA, Bell JT, Marioni JC, Pritchard JK, Gilad Y. 2011. A genome-wide study of DNA methylation patterns and gene expression levels in multiple human and chimpanzee tissues. PLoS genetics 7(2): e1001316.

Pai AA, Cain CE, Mizrahi-Man O, De Leon S, Lewellen N, Veyrieras JB, Degner JF, Gaffney DJ, Pickrell JK, Stephens M et al. 2012. The contribution of RNA decay quantitative trait Loci to inter-individual variation in steady-state gene expression levels. PLoS genetics 8(10): e1003000.

Pique-Regi R, Degner JF, Pai AA, Gaffney DJ, Gilad Y, Pritchard JK. 2011. Accurate inference of transcription factor binding from DNA sequence and chromatin accessibility data. Genome research 21(3): 447–455.

Pollard KS, Hubisz MJ, Rosenbloom KR, Siepel A. 2010. Detection of nonneutral substitution rates on mammalian phylogenies. Genome Res 20(1): 110–121.

Pollard KS, Salama SR, Lambert N, Lambot MA, Coppens S, Pedersen JS, Katzman S, King B, Onodera C, Siepel A et al. 2006. An RNA gene expressed during cortical development evolved rapidly in humans. Nature 443(7108): 167–172.

Prabhakar S, Visel A, Akiyama JA, Shoukry M, Lewis KD, Holt A, Plajzer-Frick I, Morrison H, Fitzpatrick DR, Afzal V et al. 2008. Human-specific gain of function in a developmental enhancer. Science 321(5894): 1346–1350.

Rada-Iglesias A, Bajpai R, Swigut T, Brugmann SA, Flynn RA, Wysocka J. 2011. A unique chromatin signature uncovers early developmental enhancers in humans. Nature 470(7333): 279–283.

Redon R, Ishikawa S, Fitch KR, Feuk L, Perry GH, Andrews TD, Fiegler H, Shapero MH, Carson AR, Chen W et al. 2006. Global variation in copy number in the human genome. Nature 444(7118): 444–454.

Robertson AG, Bilenky M, Tam A, Zhao Y, Zeng T, Thiessen N, Cezard T, Fejes AP, Wederell ED, Cullum R et al. 2008. Genome-wide relationship between histone H3 lysine 4 mono- and tri-methylation and transcription factor binding. Genome research 18(12): 1906–1917.

Rockman MV, Hahn MW, Soranzo N, Zimprich F, Goldstein DB, Wray GA. 2005. Ancient and recent positive selection transformed opioid cis-regulation in humans. PLoS Biol 3(12): e387.

Ruthenburg AJ, Allis CD, Wysocka J. 2007. Methylation of lysine 4 on histone H3: intricacy of writing and reading a single epigenetic mark. Mol Cell 25(1): 15–30.

Rybtsova N, Leimgruber E, Seguin-Estevez Q, Dunand-Sauthier I, Krawczyk M, Reith W. 2007. Transcription-coupled deposition of histone modifications during MHC class II gene activation. Nucleic acids research 35(10): 3431–3441.

Santos-Rosa H, Schneider R, Bannister AJ, Sherriff J, Bernstein BE, Emre NC, Schreiber SL, Mellor J, Kouzarides T. 2002. Active genes are tri-methylated at K4 of histone H3. Nature 419(6905): 407–411.

Santos-Rosa H, Schneider R, Bernstein BE, Karabetsou N, Morillon A, Weise C, Schreiber SL, Mellor J, Kouzarides T. 2003. Methylation of histone H3 K4 mediates association of the Isw1p ATPase with chromatin. Mol Cell 12(5): 1325–1332.

Schones DE, Cui K, Cuddapah S, Roh TY, Barski A, Wang Z, Wei G, Zhao K. 2008. Dynamic regulation of nucleosome positioning in the human genome. Cell 132(5): 887–898.

Shapiro MD, Marks ME, Peichel CL, Blackman BK, Nereng KS, Jonsson B, Schluter D, Kingsley DM. 2004. Genetic and developmental basis of evolutionary pelvic reduction in threespine sticklebacks. Nature 428(6984): 717–723.

Shulha HP, Crisci JL, Reshetov D, Tushir JS, Cheung I, Bharadwaj R, Chou HJ, Houston IB, Peter CJ, Mitchell AC et al. 2012. Human-specific histone methylation signatures at transcription start sites in prefrontal neurons. PLoS biology 10(11): e1001427.

Siepel A, Bejerano G, Pedersen JS, Hinrichs AS, Hou M, Rosenbloom K, Clawson H, Spieth J, Hillier LW, Richards S et al. 2005. Evolutionarily conserved elements in vertebrate, insect, worm, and yeast genomes. Genome Res 15(8): 1034–1050.

Song Q, Smith AD. 2011. Identifying dispersed epigenomic domains from ChIP-Seq data. Bioinformatics 27(6): 870–871.

Stranger BE, Forrest MS, Clark AG, Minichiello MJ, Deutsch S, Lyle R, Hunt S, Kahl B, Antonarakis SE, Tavare S et al. 2005. Genome-wide associations of gene expression variation in humans. PLoS genetics 1(6): e78.

Stranger BE, Nica AC, Forrest MS, Dimas A, Bird CP, Beazley C, Ingle CE, Dunning M, Flicek P, Koller D et al. 2007. Population genomics of human gene expression. Nature genetics 39(10): 1217–1224.

Tippmann SC, Ivanek R, Gaidatzis D, Scholer A, Hoerner L, van Nimwegen E, Stadler PF, Stadler MB, Schubeler D. 2012. Chromatin measurements reveal contributions of synthesis and decay to steady-state mRNA levels. Molecular systems biology 8: 593.

Verlaan DJ, Ge B, Grundberg E, Hoberman R, Lam KC, Koka V, Dias J, Gurd S, Martin NW, Mallmin H et al. 2009. Targeted screening of cis-regulatory variation in human haplotypes. Genome research 19(1): 118–127.

Veyrieras JB, Kudaravalli S, Kim SY, Dermitzakis ET, Gilad Y, Stephens M, Pritchard JK. 2008. High-resolution mapping of expression-QTLs yields insight into human gene regulation. PLoS Genet 4(10): e1000214.

Wang Z, Zang C, Rosenfeld JA, Schones DE, Barski A, Cuddapah S, Cui K, Roh TY, Peng W, Zhang MQ et al. 2008. Combinatorial patterns of histone acetylations and methylations in the human genome. Nature genetics 40(7): 897–903.

Warner LR, Babbitt CC, Primus AE, Severson TF, Haygood R, Wray GA. 2009. Functional consequences of genetic variation in primates on tyrosine hydroxylase (TH) expression in vitro. Brain Res 1288:1–8.

Woychik NA, Hampsey M. 2002. The RNA polymerase II machinery: structure illuminates function. Cell 108(4): 453–463.

Wray GA, Hahn MW, Abouheif E, Balhoff JP, Pizer M, Rockman MV, Romano LA. 2003. The evolution of transcriptional regulation in eukaryotes. Mol Biol Evol 20(9): 1377–1419.

Xiao S, Xie D, Cao X, Yu P, Xing X, Chen CC, Musselman M, Xie M, West FD, Lewin HA et al. 2012. Comparative epigenomic annotation of regulatory DNA. Cell 149(6): 1381–1392.

Zeitlinger J, Stark A, Kellis M, Hong JW, Nechaev S, Adelman K, Levine M, Young RA. 2007. RNA polymerase stalling at developmental control genes in the Drosophila melanogaster embryo. Nature genetics 39(12): 1512–1516.

Zeller T, Wild P, Szymczak S, Rotival M, Schillert A, Castagne R, Maouche S, Germain M, Lackner K, Rossmann H et al. 2010. Genetics and beyond—the transcriptome of human monocytes and disease susceptibility. PLoS One 5(5): e10693.

Zhang Y, Liu T, Meyer CA, Eeckhoute J, Johnson DS, Bernstein BE, Nussbaum C, Myers RM, Brown M, Li W et al. 2008. Model-based analysis of ChIP-Seq (MACS). Genome biology 9(9): R137.

Zhou X, Stephens M. 2012. Polygenic modeling with Bayesian sparse linear mixed models.

